# Modelling perception as a hierarchical competition differentiates imagined, veridical, and hallucinated percepts

**DOI:** 10.1101/2022.09.02.506121

**Authors:** Alexander A Sulfaro, Amanda K Robinson, Thomas A Carlson

**Affiliations:** School of Psychology, The University of Sydney, Sydney, New South Wales, Australia; Queensland Brain Institute, The University of Queensland, Brisbane, Queensland, Australia

**Keywords:** mental imagery, hallucinations, aphantasia, hyperphantasia, daydreaming, perception

## Abstract

Mental imagery is a process by which thoughts become experienced with sensory characteristics. Yet, it is not clear why mental images appear diminished compared to veridical images, nor how mental images are phenomenologically distinct from hallucinations, another type of non-veridical sensory experience. Current evidence suggests that imagination and veridical perception share neural resources. If so, we argue that considering how neural representations of externally-generated stimuli (i.e. sensory input) and internally-generated stimuli (i.e. thoughts) might interfere with one another can sufficiently differentiate veridical, imaginary, and hallucinatory perception. We here use a simple computational model of a serially-connected, hierarchical network with bidirectional information flow to emulate the primate visual system. We show that modelling even first-approximations of neural competition can more coherently explain imagery phenomenology than non-competitive models. Our simulations predict that, without competing sensory input, imagined stimuli should ubiquitously dominate hierarchical representations. However, with competition, imagination should dominate high-level representations but largely fail to outcompete sensory inputs at lower processing levels. To interpret our findings, we assume low-level stimulus information (e.g. in early visual cortex) contributes most to the sensory aspects of perceptual experience, while high-level stimulus information (e.g. towards temporal regions) contributes most to its abstract aspects. Our findings therefore suggest that ongoing bottom-up inputs during waking life may prevent imagination from overriding veridical sensory experience. In contrast, internally-generated stimuli may be hallucinated when sensory input is dampened or eradicated. Our approach can explain individual differences in imagery, along with aspects of daydreaming, hallucinations, and non-visual mental imagery.

## Introduction

Mental images are internally-generated thoughts which can be seen, heard, or in some way perceived with sensory qualities. Yet, how does the brain generate sensory experiences without a real sensory stimulus in the environment to perceive? One answer might be that mental imagery involves similar processes to those used during veridical perception, yet with an opposite direction of information flow. Instead of pooling together sensory information to extract abstract knowledge from a real stimulus, mental imagery may involve retrohierarchically reconstructing the sensory features of an imagined stimulus from abstract knowledge we already possess. In line with this idea, mental imagery seems to use the same neural machinery as that used during veridical perception (Cichy et al., 2012; Dijkstra et al., 2017, 2018; Dijkstra, Bosch, et al., 2019; Ishai, 2010; Kosslyn et al., 1993; Xie et al., 2020), yet with neural activity during the early stages of veridical visual perception resembling neural activity during the later stages of visual mental imagery and vice versa (Breedlove et al., 2020; Dentico et al., 2014; Dijkstra et al., 2020; Linde-Domingo et al., 2019).

However, mental imagery is unlikely to be an exact reverse of feedforward processes in veridical perception given that individuals reliably report that their mental images are experienced with some form of reduced sensory quality (termed *vividness*) compared to real images (Galton, 1880; Marks, 1973). Some researchers have proposed that this difference follows from differences between feedforward and feedback processes, the latter of which are believed to be responsible for propagating mental images. For instance, information about mental imagery content tends to be most detectable in superficial and deep cortical layers, but not granular mid-layers, in contrast to information about environmental stimulus content which can be detected across all cortical layers (Bergmann et al., 2019; Lawrence et al., 2018). Other accounts speculate that imagery could be propagated using weak feedback connections in the visual system, in contrast to feedforward connections which stereotypically drive, rather than modulate, action potentials (Koenig-Robert & Pearson, 2021). However, feedback to early visual regions can drive action potentials depending on local neurochemistry (Aru et al., 2020). Further, neither of these differences between feedforward and feedback processes can explain the unique experience of mental imagery without a framework specifying what it means for a mental image to be a quasi-sensory experience to begin with.

A more readily-interpretable explanation is that mental images appear impoverished simply because the features of the images they depict are distorted relative to those of real images. That is, mental images might merely have less-precise image statistics (e.g. spatial frequency, location, size) compared to real images, but are otherwise just as visible. Breedlove et al. (2020) provide an account which allows for this possibility, finding that representations of mental images in early visual cortices are deficient in detailed sensory information compared to real images. As hierarchical feedforward processes combine multiple pieces of sensory information (e.g. edges) to form fewer pieces of more abstract information (e.g. a shape), reconstructing sensory information using retrohierarchical feedback requires extrapolating from a relatively information-deficient source material. This process should distort imagined stimuli in predictable ways such that neural populations which represent mental images in early visual cortices have lower spatial frequency preferences, more foveal receptive fields, and larger receptive fields than when responding to real images (Breedlove et al., 2020).

Ultimately, many factors may distinguish between veridical and mental percepts given that the former are externally-generated while the latter are internally-generated. However, any complete explanation of the unique appearance of mental images must also account for how the quasi-sensory experience of mental imagery might be differentiated from other internally-generated, yet unambiguously-sensory experiences, such as which occur in hallucinations, many dreams, and eidetic mental imagery. Regardless of the image statistics of the internally-generated content being depicted, these latter phenomena tend to involve a sensory experience of internally-generated content which is entirely equivalent to the subjective experience of externally-generated content (Chiu, 1989; Ffytche et al., 1998) except that the content experienced is non-veridical and is not typically experienced as part of normal waking life. For the purposes of this article, we refer to the internally-generated, unambiguously-sensory experience common to each of hallucinations, dreams, and eidetic imagery as hallucinatory imagery, or a hallucination generally, regardless of whether such imagery is voluntary or involuntary, regardless of whether it is believed to be real or unreal, and regardless of where exactly within the visual field it is perceived to be located. Crucially, while mental imagery is quasi-sensory, seemingly seen yet unseen, hallucinatory imagery is definitively visible. This should require that non-veridical internally-generated sensory content or features replace veridical sensory content or features, overwriting them in a given region of the visual field. For instance, seeing a hallucinatory cat on this page instead of text, or seeing the text on this page as moving instead of still, or seeing anything at all during a dream instead of merely the back of our eyelids.

Here, we propose that imagined, veridical, and hallucinated percepts can be clearly distinguished by considering perception as a competitive process. While mental imagery generation can be modelled without competition from veridical perception (e.g. Breedlove et al., 2020), mental imagery almost never occurs in isolation. Instead, mental images are invariably generated while we are awake and receiving sensory input from the external world. If mental imagery and veridical perception share neural resources, then both processes should interact such that neural representations of imagined and real stimuli interfere or compete with one another.

Interactions between real and imagined stimuli have been previously demonstrated, lending support to the idea that perception may be a competitive process, although the exact outcome and nature of this interaction seems to vary. For instance, mental images can constructively interfere with real image content, depending on stimulus congruency (Dijkstra et al., 2022), and can induce priming or adaptation to veridical stimuli, depending on the individual (Dijkstra et al., 2021; Dijkstra, Hinne, et al., 2019; Keogh & Pearson, 2017). Increasing environmental luminance can also impede the priming effects of mental imagery (Keogh & Pearson, 2017), while real modality-specific distractors reduce the recall of modality-specific information (Vredeveldt et al., 2011; Wais et al., 2010), potentially reflecting competitive interference.

We sought to predict whether perceptual interference could account for the quasi-sensory, rather than hallucinatory, experience of mental imagery. Yet, any valid explanation of the quasi-sensory experience of mental imagery must first clearly specify what it means to have a quasi-sensory experience at all. Such an experience may be possible under a framework where perceptual experience itself is considered as a multifaceted phenomenon, composed of both sensory and abstract experiences at any given time. In vision, this entails having an experience of the low-level features of an image, such as its hues and edge orientations, while also having an experience of the high-level features of what the image actually depicts, such as recognising that an image is depicting a duck. Usually, low- and high-level features of an image seem intertwined such that it is unintuitive to consider them as separate components of perceptual experience. Yet, these aspects become clearly separable when we consider bistable percepts (e.g. the rabbit-duck illusion), where the same low-level information can give rise to different abstract experiences. These abstract experiences feel perceptually distinct from one another even though the spatial pattern of light in our visual field, and our purely low-level visual experience of it, remains unchanged.

To anchor these experiences to physical processes, we assume that the sensory and abstract aspects of experience are dependent on neural systems encoding each respective type of information in the brain, such as those along the visual ventral stream. However, as neural representations of externally- and internally-generated stimuli are not explicitly sensory or abstract, but distributed on a spectrum, so too may perceptual experience itself be composed of a spectrum of sensory and abstract aspects. Given that internally-generated perceptual experiences also seem to use similar neural resources to their externally-generated counterparts, including in a feature-specific manner (Ffytche et al., 1998), then mental and hallucinatory imagery processes may also be experienced with a spectrum of abstract and sensory features. Under this framework, the quasi-sensory experience of mental imagery can be clearly conceptualised as the experience arising from the activation of high-level to mid-level, but not low-level, feature representations of internally-generated content, regardless of the content itself. Hallucinatory imagery, in contrast, may invoke low-level sensory representations, with or without more abstract components, whilst veridical percepts should at least invoke low-level featural representations given their bottom-up entryway into the visual system.

Under these assumptions, we constructed simple computational model of a serially-connected network with bidirectional information flow as a first-approximation to simulating the hierarchical interaction of internally- and externally-generated stimulus representations (Figure 1). We used this model to evaluate the degree to which internally-generated (i.e. imagined) stimulus representations are able to spread throughout a hierarchical system, with and without competition from externally-generated stimuli. We demonstrate that competition from external sensory input could prevent internally-generated content from ever recruiting the low-level neural infrastructure best suited to representing highly-sensory, highly-modal information. In doing so, we show that accounting for perceptual interference, even with preliminary models, can provide an arguably more coherent explanation for the unique phenomenology of mental imagery than approaches which consider mental imagery in isolation. We explore the perceptual implications of such predictions on the subjective experience of mental imagery, provide a neurochemically-grounded hypothesis for variance in imagery across individuals and states, and delineate how perceptual interference may relate to hallucinations and mental imagery in non-visual modalities. Ultimately, we demonstrate that reconceptualising mental imagery as an intrinsically competitive process can resolve major lingering questions in imagery research.

**Figure 1.**
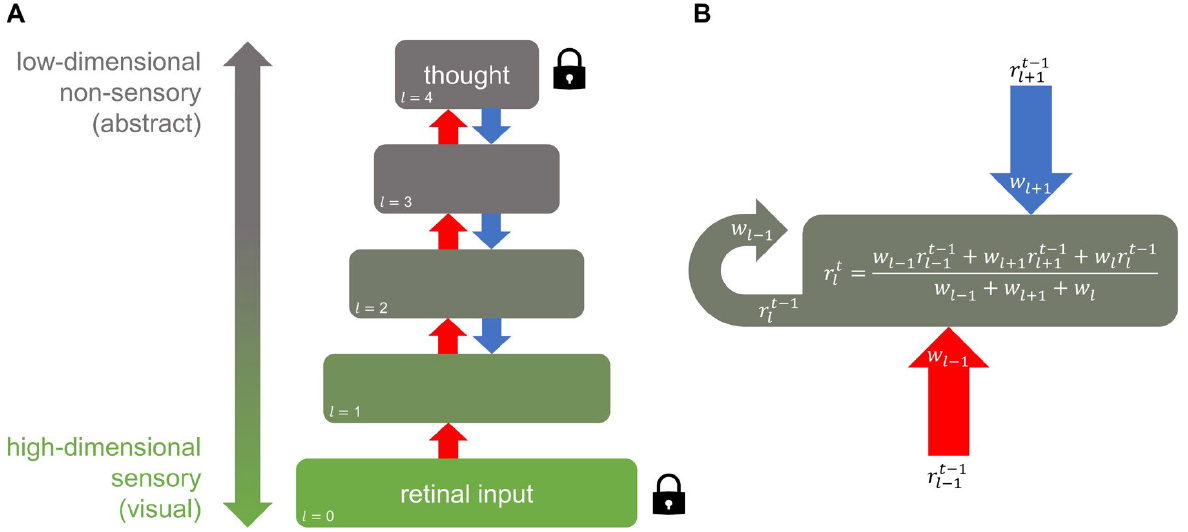
A hierarchical network modelling interference between externally- and internally-generated stimuli. *Note*. **A**. A hierarchical network model of visual perception. Layer size indicates the relative dimensionality of the encoded representation. Information is pooled via feedforward processes (red) and extrapolated via feedback processes (blue). Lock icons indicate input layers with clamped representations during simulations where mental imagery is attempted during veridical perception. Clamped representations do not evolve with time. Note that a “thought” here could be any internally-generated representation. **B**. Evolution rule for unclamped layers. Each value *r* in the representational matrix of each layer *l* evolves at each timepoint *t* according to a weighted average of the adjacent layers and the current layer at the previous timestep with weights *w*. The ascending value array was convolved with an averaging kernel before being weighted to facilitate feedforward dimensionality reduction.

## Methods

### Computational model

Our goal was to assess the degree and extent to which internally-generated stimuli are able to spread throughout a hierarchical system, with and without competition from externally-generated stimuli. To do this, a dynamic, serially-connected, five-layer computational network was constructed as a simple model of how stimulus information may flow bidirectionally in the hierarchical human visual system (Figure 1). Each layer *l* encoded a representation *r* consisting of a square matrix of values. The number of units available to represent information (i.e. dimensionality) in each layer decreased linearly in size from 256 × 256 matrix elements at the lowest layer (*l = 0*) to 208 × 208 elements at the higher layer (*l = 4*). This mimics feedforward dimensionality reduction in the visual system such that upper layers encode more abstract stimulus information than lower layers, analogous to neural structures towards the parietal and temporal lobes (Binder, 2016). Each element in the matrix could vary in value along a single dimension, from 0 to 1, encoding the variation of an arbitrary stimulus feature. The lowest layer acted as an input layer for externally-generated stimuli and never received feedback, analogous to the retina in visual perception.

The network allowed representations to propagate between layers, and for bottom-up processes to interact with top-down processes. Hence, the representation *r* of each layer *l* was updated at each timestep *t* according to a weighted average of the ascending representation *r*_*l-1*_ from the layer below, descending representation *r*_*l+1*_ from the layer above, and representation *r*_*l*_ of the current layer, each at the previous timestep *t-1* (Figure 1B). We note that despite the simplicity of this approach, weighted averaging is approximately equivalent to the biologically-plausible additive combination of neural inputs, followed by a decay of activity proportional to the strength of activation. However, unlike in real neurons, decay here occurs instantaneously. We allow for this simplification given that we simulate the perception of static images only and are therefore most concerned with the steady-state, rather than moment-to-moment, profile of the system.

Each layer was updated sequentially, first as part of a feedforward sweep, then as part of a feedback sweep, and then alternating with every subsequent iteration. Weightings *w*_*l-1*_, *w*_*l+1*_, and *w*_*l*_ for the ascending, descending, and current representations respectively were each equal to one except where the effect of weighting ratios was explicitly investigated. However, if a layer acted as a constant source of information in the network (representing either an internally-or externally-generated stimulus), the values in its representational matrix were locked in place (clamped) by setting the weightings of ascending and/or descending inputs to that layer to zero. Given that the retina is always receiving visual input in an awake state, the lowest layer of the network was always clamped to some predetermined set of input values. Drawing from modelling by Breedlove et al. (2020), the highest layer in the network was also clamped to a fixed set of input values to simulate an internally-generated stimulus (i.e. a thought) during mental imagery.

To simulate feedforward information pooling in the visual ventral stream, ascending representations were combined using a 13 × 13 square convolutional kernel such that each element of the layer above received the average of inputs from multiple elements from the layer below. Hence, from *l = 0* to *l = 4*, the receptive field size of each element in each representation increased while layer dimensionality decreased. Because forming a mental image of an object first requires at some point seeing the object or its constituent parts, internally-generated stimulus representations were formed from externally-generated (e.g. retinal) stimulus inputs first processed through a single complete feedforward sweep from layer 0 to layer 4.

To assess the degree to which mental imagery was able to compete with veridical sensory input, we allowed the network to evolve under three main sets of initial conditions: veridical perception without imagination, imagination without veridical perception, and imagination during veridical perception. In the first case, to simulate a scenario where externally-generated visual input is present without competition, only the bottom layer was clamped to a stimulus representation (veridical perception without imagination). Secondly, to simulate a scenario where an internally-generated stimulus is present without competition, the top layer was clamped to an imagined stimulus representation (imagination without veridical perception). The ascending input from the base layer was down-weighted to zero in this scenario to enforce a state of sensory disconnection. Thirdly, in what we consider a more realistic mental imagery scenario, both the top and bottom layers were simultaneously clamped to different stimulus representations to simulate mental imagery occurring in the presence of competing sensory input (imagination during veridical perception). All intervening or non-input layers were initialised with white noise.

In each scenario, we measured the steady-state contribution of top-down representations relative to bottom-up representations at each layer. This was achieved by taking the average value of all elements in each layer for the case where the initial low-level or high-level input representations were set homogenously to zeros or ones, respectively (Figure 2D-F). Each scenario was also replicated using images as inputs with each element of a representation corresponding to one pixel in the image such that the collective pixel values of each image equate to a simulated neural activity pattern (Figure 2A-C). A 256 × 256 greyscale image of a camel was used for sensory input, and a 256 × 256 greyscale image of a hat was used for imagined input after conversion to a reduced dimensionality of 208 × 208 via feedforward pooling. Both were sourced from the BOSS image database (Brodeur et al., 2010). The similarity between layer activity patterns and the original imagined or veridical stimulus patterns can be quantified by calculating the absolute mean pixel difference between the images. We also separately explored how varying the ascending and descending weighting schemes affected competition between simulated imagined and veridical stimulus representations (Figure 3). For each scenario in Figure 3, all mid-layer feedforward or feedback weightings were ≤ 1 with the ratio specified in the figure for the duration of the simulation.

**Figure 2.**
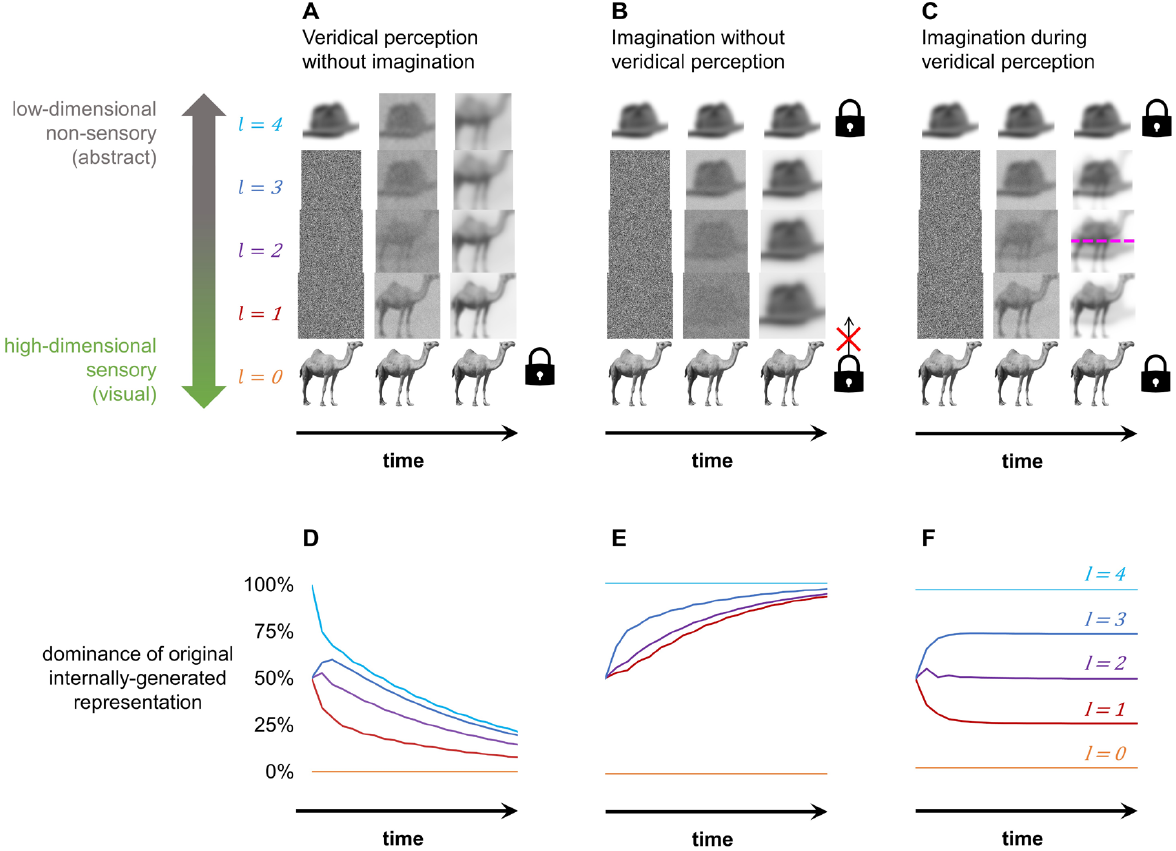
Veridical stimuli outcompete imagined stimuli at low-level sensory representations, but not high-level abstract representations. *Note*. How simulated neural representations of an internally-generated (imagined) hat and an externally-generated (veridical) camel spread through a perceptual processing hierarchy with and without competition. **A-C**. Simulated neural activity patterns, coded as image pixel values, evolving over 0, 2, and 20 alternating feedforward or feedback sweeps. Initial activity patterns are the same in each scenario, and mimic thinking about a hat while receiving a retinal image of a camel. Lock icons indicate clamped representations which do not evolve. The crossed arrow indicates disconnected sensory input (w_l = 0_ = 0). **A**. All layers except the base layer are free to evolve, causing activity patterns to be dominated by the initial veridical input. **B**. Mid-layers are free to evolve only. Sensory input is present but disconnected. All evolving activity patterns become dominated by the initial imagined input. **C**. Mid-layers are free to evolve only, and sensory input is present and connected. Imagined content dominates upper layers and veridical content dominates lower layers. The magenta line indicates the *representational equilibrium point* where the internally- and externally-generated stimuli equally contribute to a layer. **D-F**. One-dimensional simulations of each scenario above in A-C illustrating how the character of each layer evolves over time. The dominance of the original externally-generated stimulus representation at *l = 0* can be obtained by subtracting the internally-generated stimulus dominance from 100%.

**Figure 3.**
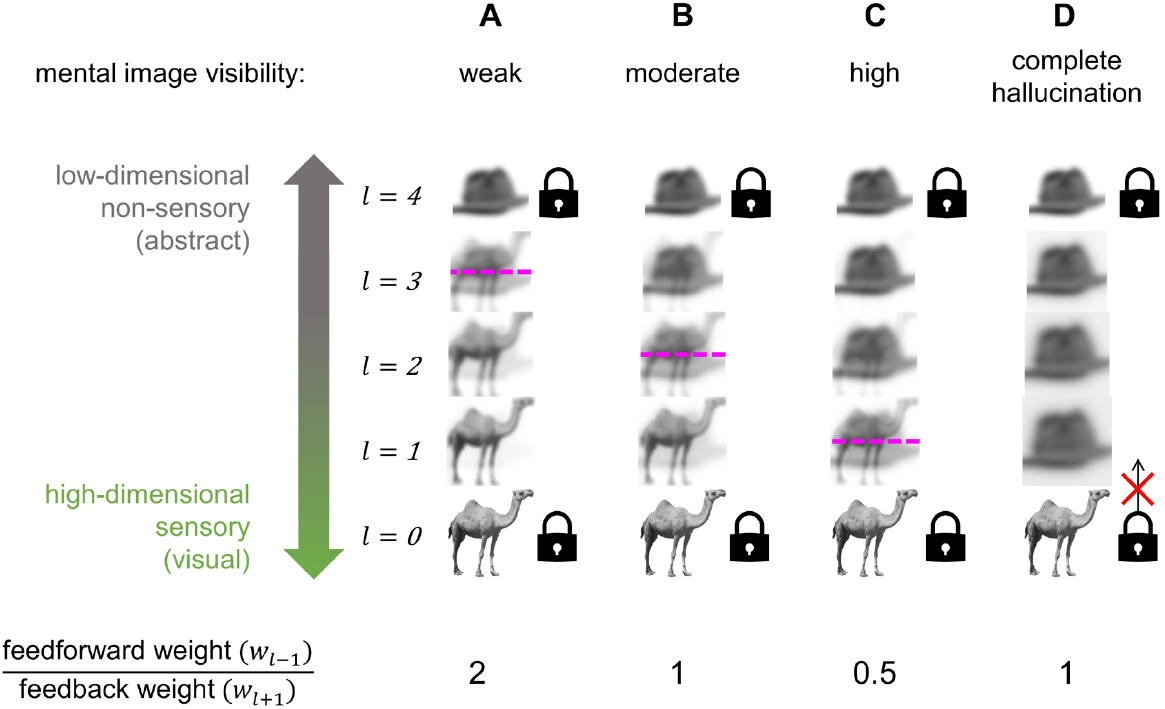
Decreasing feedforward-to-feedback weighting ratio induces more sensory mental imagery. *Note*. Hierarchical competition between simulated neural representations of an internally-generated (imagined) hat and an externally-generated (veridical) camel modelled under different feedforward-to-feedback weighting ratios. Simulated neural activity patterns, coded as image pixel values, are depicted after 20 alternating feedforward or feedback sweeps. Initial activity patterns are the same in each scenario, and mimic thinking about a hat while receiving a retinal image of a camel. Lock icons indicate clamped representations which do not evolve. The crossed arrow indicates disconnected sensory input (w_l = 0_ = 0). The magenta line indicates the *representational equilibrium point* where the internally- and externally-generated stimuli contribute equally to a layer. **A-C**. Globally varying the ratio of bottom-up (*w*_*l-1*_) and top-down (*w*_*l+1*_) weightings affects representational equilibrium and the extent to which thoughts should be experienced visually (e.g. weak, moderate, or high). **D**. All unclamped layers converge towards the original internally-generated stimulus representation once sensory input is disconnected.

### Model interpretation

Our model can be interpreted as analogous to a simplified model of the human visual processing hierarchy, with a basal layer encoding externally-generated retinal input and an apical layer housing more abstract internally-generated stimulus representations encoded by neural structures towards the parietal and temporal lobes (Binder, 2016). To interpret simulation outcomes phenomenologically, we considered that a single perceptual experience can contain a distribution of information, spanning from highly-sensory (modal) in nature to highly-abstract (amodal). We assume that each representational layer in the visual hierarchy which receives both feedforward and feedback input contributes information to the construction of conscious perceptual experience. We also assume that the type of information provided by each layer aligns with the layer’s dimensionality such that high-dimensional representations contribute most to the sensory aspects of the experience while low-dimensional representations contribute most to the non-sensory aspects of the experience. Hence, low-level representations would be most influential in determining the literal image content of a visual experience (e.g. the exact location, orientation, and contrast of high-spatial frequency edges), while high-level representations would contribute most to the abstract understanding of what is being thought about or observed (e.g. knowing whether the edges in a scene form a hat or a camel), with a spectrum of quasi-sensory or quasi-abstract features in between.

Crucially, this implies that a perceptual experience *gains* sensory or abstract properties depending on how well neural regions with the corresponding high-or low-dimensional structure, respectively, are recruited during that experience. Therefore, the further an internally-generated stimulus representation is able to spread towards the primary visual cortex, for instance, the more it may be experienced with properties approaching that of an actual visual image (i.e. a true mental image) rather than something pondered without sensory qualia (i.e. a non-sensory thought). This approach can therefore maintain consistency with traditional accounts implicating the primary visual cortex in mental imagery (Kosslyn, 2005; Kosslyn & Thompson, 2003; Pearson, 2019; Pearson & Kosslyn, 2015).

Note that, given hallucinatory visual experience is possible without corresponding retinal activity, we consider the lowest level in the hierarchy purely as a non-conscious input layer to the rest of the network. We also note that while there’s compelling evidence that some high-level representations are stored in a largely amodal form (e.g. Popham et al., 2021), they are not necessarily completely non-sensory. Late-stage object representations can still contain some degree of sensory information (e.g. see DiCarlo & Cox, 2007), albeit in a considerably more low-dimensional form relative to primary sensory areas. Our model only requires higher-level representations to be less sensory than low-level representations rather than completely non-sensory.

## Results

### Perception without competition

Two scenarios were simulated where competition was not present: veridical perception without imagination, and imagination without veridical perception. Initiation of veridical perception was simulated by allowing the lowest representational layer to have a non-zero contribution to the layer above. Imagination was simulated by preventing the highest layer from being affected by feedforward inputs below such that it provided a constant source of top-down information to the system. In both cases, only one fixed source of perceptual information was present within the network, either an externally-or internally-generated stimulus representation. An example simulation is shown in Figure 2 using an image of a camel as the externally-generated veridical stimulus input (i.e. retinal activity pattern), and a compressed image of a hat as the internally-generated imagined stimulus input (i.e. high-level neural activity pattern).

Without competition from imagination, all freely-evolving (unclamped) layers in the network converged asymptotically to a representation dominated by the externally-generated stimulus representation (Figure 2A). Conversely, without competition from veridical perception, all unclamped layers converged to a representation dominated by the internally-generated stimulus representation (Figure 2B). As all unclamped representations are eventually dominated by the original externally-or internally-generated source stimulus, both scenarios each result in a state corresponding to having a visual experience of the same stimulus which is being thought about abstractly. In the case of pure veridical perception, the system evolves to emulate a common perceptual experience: seeing an object while also knowing what it is. Yet, where the internally-generated stimulus is uncontested, the sensory characteristics of the imagined stimulus should completely dominate visual experience, resulting in the perception of what was imagined and not what was present in the environment. Hence, pure imagination imitates hallucinating the object one is thinking about.

Although in both cases, layer *l = 1* is entirely dominated by one stimulus, each state occurred through a distinct formation route. Note that the top-down, but not bottom-up, formation route resulted in substantial distortions at lower layers, reflecting the findings of Breedlove et al. (2020). Yet, these distortions only affect the *content* of the imagined stimulus, not the degree to which that stimulus is able to commandeer a hierarchical layer in this model.

### Perception with competing imagined and veridical stimuli

Where two fixed sources of information are simultaneously present within the network, neither the imagined nor veridical stimulus can dominate every unclamped layer in the system (Figure 2C). Instead, the network represents both stimuli simultaneously at the steady state. However, the further a given layer is from an externally-or internally-generated stimulus input layer, the smaller the contribution of the original veridical or imagined stimulus to the representation held by the given layer. This is shown in Figure 2F for the case where feedforward and feedback weightings are equal (wl-1 = w_l+1_ = 1).

Consequently, where imagination occurs in the presence of sensory input, the externally-generated veridical stimulus dominates low-level representations while the internally-generated imagined stimulus dominates high-level representations. Such a system is therefore representing two distinct stimuli, yet at largely separate hierarchical levels. As lower levels are more suited for representing sensory information, and higher levels for abstract, this state corresponds to a situation where an individual has a visual experience of their veridical environment despite simultaneously having relatively non-sensory thoughts about what they are imagining. Hence, the simulation suggests that external sensory input outcompetes imagination at the low-level regions most likely to contribute to the most sensory aspects of perceptual experience.

### Modelling variation in imagery quality (e.g. aphantasia and hyperphantasia) as a shift in representational equilibrium

This model can also be used to understand states where thoughts are experienced in a highly sensory way (hyperphantasia) or in a non-sensory or weakly-sensory way (aphantasia; Figure 3). When sensory input and imagination compete, our simulation showed that spanning along the perceptual hierarchy from bottom to top corresponded to transitioning from primarily representing a veridical stimulus to primarily representing an imagined stimulus. The exact point where the two stimuli are represented in equal proportion is here dubbed the *representational equilibrium point* for the system. Representations contain more veridical stimulus character below this point and more imagined stimulus character above it.

The location of the representational equilibrium point can summarise how far an imagined representation can penetrate towards lower layers before it is outcompeted by the veridical stimulus, or how far a veridical stimulus representation can penetrate towards upper layers before it is outcompeted by the imagined stimulus. Where ascending and descending information is weighted equally (*w*_*l-1*_ = *w*_*l+1*_ = 1; Figure 3B), the equilibrium point lies at the middle layer in the centre of the hierarchy such that lower layers (e.g. towards the primary visual cortex) are dominated by externally-generated sensory input and upper layers (e.g. towards the temporal lobe) are dominated by internally-generated thoughts, symmetrically. However, overweighting feedforward information (*w*_*l-1*_/*w*_*l+1*_ > 1) elevates the equilibrium point such that the imagined stimulus is restricted to only the highest, least-modal hierarchical layers, corresponding to a state of aphantasia (Figure 3A). In contrast, overweighting feedback information (*w*_*l-1*_/*w*_*l+1*_ < 1) lowers the equilibrium point such that the imagined stimulus dominates the majority of the hierarchy, corresponding to a state of hyperphantasia (Figure 3C). Feedforward-to-feedback weighting ratio variation could therefore account for between-person and moment-to-moment differences in the apparent sensory quality of mental imagery experiences.

## Discussion

### Distinguishing imagined, veridical, and hallucinatory percepts

In this study, we aimed to explain the apparent differences in vividness between veridical, imagined, and hallucinated percepts. We show that, for a hierarchical system with bidirectional information flow, the presence of competing sensory input can prevent internally-generated stimuli from dominating low-level regions most suited for supporting a modal, highly-sensory neural representation. We interpret this finding as suggesting that, when competing sensory input is present, the sensory content of imagination does not supersede the sensory content of veridical perception. That is, for instance, even if we imagine a hat, our mental image of the hat does not replace the image of a real camel in our visual field: the camel is still more visible. In contrast, the attenuation or abolition of competing external sensory input could facilitate internally-generated stimuli becoming the dominant contributor to sensory experience, creating hallucinatory imagery. Our model therefore illustrates that the apparent difference in reported vividness between veridical and imagined percepts could be explained by the degree to which imagined and veridical stimulus representations spread hierarchically, and that this difference can arise due to external sensory input competing with our internally-generated thoughts.

### Neurological consequences of perceptual competition

Our model results in some counterintuitive implications. Despite the suitability of the early visual cortex for representing retinotopic, visual information, our account suggests that hierarchically low-level neural regions (e.g. primary sensory cortices) may be less involved in representing an imagined stimulus than any other region in the corresponding perceptual hierarchy. However, this implication is consistent with existing findings. Individuals still report being able to create mental images despite near-complete bilateral lesions to primary visual cortex, suggesting that the primary visual cortex is not necessary for visual mental imagery (Bridge et al., 2012; Chatterjee & Southwood, 1995; de Gelder et al., 2015; Zago et al., 2010). Even mid-level impairments to the visual system, a potential cause of visual agnosia, can leave visual mental imagery preserved (Behrmann et al., 1994). Further, a meta-analysis of the neural correlates of visual mental imagery by Spagna et al. (2021) found that the left fusiform gyrus, a high-level region within the visual ventral stream, was reliably involved during mental imagery across studies while early visual cortices were not.

These findings are entirely in line with the predictions of our model: while primary sensory cortices can be involved in mental imagery, the only essential component of the experience is the internal generation of a non-veridical stimulus representation, which in our model occurs at the hierarchical apex, corresponding, for instance, to late in the visual ventral stream. This may be why subjective vividness ratings better correlate to how distinctly imagined information is represented in retinotopic, rather than associative, neural regions (S.-H. Lee et al., 2012). Under our model, activation at levels below the clamped internally-generated representation can add progressively more sensory components to the experience of imagining, although at what stage a relatively amodal, abstract thought formally becomes a truly sensory mental image may be an arbitrary distinction.

Our simulations predict that the dominance of internally-generated stimuli should increase from posterior to anterior regions of the visual hierarchy. Supporting this prediction, VanRullen & Reddy (2019) used a mental imagery decoding paradigm to show that more information about imagined face images was present in temporal regions than occipital regions. Lee et al. (2012) likewise showed that less information about imagined object images could be decoded towards more posterior, retinotopic areas of the visual system. Both studies also found that externally- and internally-generated stimulus representations became more similar towards higher-level regions, aligning with the notion that imagined content is propagated from areas active during late-stage veridical perception.

### Phenomenological consequences of perceptual competition

Our model has implications for the functional and subjective characteristics of imagery. For instance, the relegation of imagination to relatively high-level representations suggests that thinking in a non-sensory way may be more commonplace than manifesting thoughts in a sensory way (e.g. as an actual visual image). This makes pragmatic sense in everyday life: thinking about what you are seeing is often favourable during veridical visual perception, yet it may be quite unfavourable to see what you are thinking about given that such hallucinations may supersede important survival-relevant environmental information. However, this process could involve a functional trade-off as the semantic content of sensory input may be processed poorly if high-level regions are already occupied by unrelated thoughts. Mental imagery could push internally-generated high-level representations towards mid-level regions, interfering with the ascending processing of externally-generated stimuli. This could explain why internal attention can affect the processing of environmental stimuli, such as during daydreaming (Schooler et al., 2011; Smallwood, 2011). States of immersive mental imagery (e.g. daydreaming) might then loosely and transiently share phenomenological similarities with agnosia given that high-level processing of externally-generated stimuli may be impeded despite low-level visual feature processing being relatively unaffected. For instance, when reading text while distracted by other thoughts, one may find themselves rereading the same paragraph repeatedly, seeing the words each time, yet failing to process the meaning of the text.

Another effect of abstract representations being activated without any significant low-level representation during imagery is that a perceiver might plausibly even *feel* like they have seen their imagined thought without ever experiencing it in any substantially visual way. That is, units involved in the recognition of an object may be activated despite minimal low-level activation. Hence people could theoretically report being able to visualise objects yet fail to be able to reconstruct the details of these imagined stimuli accurately.

Additionally, experiencing imagery on a spectrum from highly-abstract to highly-sensory may explain why the unique quasi-sensory experience of mental imagery may be difficult to describe. While the *content* of mental imagery could be described in terms of objectively-quantifiable stimulus properties (e.g. contrast, size, duration), the *experience* of such content is modified by the degree to which a mental image gains sensory features as well as just abstract features. This modulation, encapsulated by the aforementioned concept of a representational equilibrium point, could be the underlying variable commonly measured (albeit somewhat indistinctly) by subjective ratings of imagery vividness. If so, this would justify why vividness ratings are generally only investigated concerning imagined, not veridical, experiences: veridical percepts are maximally sensory by default.

### Hallucinations

Our model also delineates conditions for hallucinations. Visually, a hallucination entails seeing a non-veridical percept in a given region of the visual field *instead of* a percept that reflects external reality. Under our model, for any modality, the most perceptually compelling hallucinations would then be those where a non-veridical stimulus dominates most, if not all, low-level representations. The notion of non-veridical information penetrating sensory systems top-down to cause hallucinations has been previously explored in depth under a predictive coding framework (Powers et al., 2016). In our model, it can occur from overweighting top-down information relative to bottom-up information. However, hallucinations could also occur simply if aberrant low-level activity is fed through an otherwise normal perceptual system (Hahamy et al., 2021). Note that, in both cases, our model only accounts for the perceptual experience of imagery, hallucinated or otherwise, and not whether it was induced voluntarily (as imagery tends to be), or involuntarily (as hallucinations tend to be).

In the first possible route to hallucinations, top-down information becomes overweighted such that non-veridical thoughts can permeate low-level areas to nearly the same extent as uncontested bottom-up veridical input. In this case, hallucinations are modelled as an anomalous case of mental imagery with an extremely low representational equilibrium point, similar to hyperphantasia. If our model is correct, then individuals with hallucinatory disorders should tend to have improved mental imagery abilities. Accordingly, Parkinson’s hallucinators tend to have stronger mental imagery priming effects than non-hallucinating controls, with the degree of imagery-induced priming predicting hallucination frequency (Shine et al., 2015). Parkinson’s hallucinators also mind-wander more than non-hallucinators, corresponding to increased connectivity to the early visual cortex from default network regions (Walpola et al., 2020). Additionally, although individuals with schizophrenia can have impaired memory, their abilities on visuospatial mental imagery manipulation tasks are enhanced compared to neurotypical controls (Benson & Park, 2013; Matthews et al., 2014). However, note that some individuals are able to complete mental imagery manipulation tasks while reporting no visual mental imagery (Zeman et al., 2010).

Hallucinations should also occur when sensory input is disconnected altogether such that internally-generated percepts are entirely uncontested. If feedforward information is downweighted completely (*w*_*l-1*_ = 0) to any layer, then that layer and any layer above it are free to be commandeered by any high-level internally-generated stimulus present. Representations at such layers will then inevitably converge towards the internally-generated stimulus representation (Figure 2B, 2E, & 3D). This could contribute to hallucinations in disorders of afferent visual processing (e.g. Charles Bonnet Syndrome; Reichert et al., 2013). Hallucinations are reasonably common following cortical blindness (Aldrich et al., 1987), and in one such reported case, hallucinations could be induced directly from attempting voluntary mental imagery (Wunderlich et al., 2000). Dampening external sensory input might also contribute to hypnagogic hallucinations, which occur during sleep onset. When falling asleep, the cortex remains active several minutes after the thalamus begins silencing external sensory information (Magnin et al., 2010), potentially providing a window of time in which internally-generated cortical activity has reduced competition from the external environment.

These types of top-down influences likely aren’t the only contributor to hallucinatory perception, however. Hallucinations can be visually detailed, and sometimes reportedly even clearer than veridical percepts (Ffytche et al., 1998; Manford & Andermann, 1998; Teunisse et al., 1996). This implies hallucinations may not be a purely top-down phenomenon given that retrohierarchically extrapolating high-level information into low-level regions should result in a degradation of visual information (Breedlove et al., 2020). Many hallucinations are therefore likely to also involve aberrant low-level cortical activity (Hahamy et al., 2021; Manford & Andermann, 1998) perhaps independent of, or in tandem with, amplified feedback signals.

### Eyes-open versus eyes-closed imagery

We note that the model proposed in this paper does not presume any difference between an eyes-closed and eyes-open state apart from the change to the content of the sensory input each act produces. Though closing one’s eyes might be ostensibly interpreted as disconnecting sensory input, if this were the case, our model would predict that closing one’s eyes would immediately result in hallucinations. While hallucinations can happen when the eyes are deprived of light, such phenomena generally require either drugs (Fisher, 1991), or multiple hours (if not days) of light deprivation to manifest (Merabet et al., 2004). It could be that darkness constitutes a genuine sensory disconnection, yet with neural activity adapting to a lack of competition on slow timescales, or it could be that sensory deprivation alone does not equate to a true sensory disconnection. A formal disconnection or down-weighting of sensory input could instead require large-scale neurochemical or neurophysiological changes, such as which occur when shifting between different states of arousal. Such potential changes are described in the section below. This may be why eyes-closed hallucinations in neurotypical individuals are generally isolated to sleep onset and offset (i.e. hypnagogic and hypnopompic hallucinations) or while asleep (i.e. dreaming), but not while awake, and why hallucinations which occur in blind individuals become most likely later in the day when drowsy, regardless of light perception (Manford & Andermann, 1998).

Closing one’s eyes (at least on short time-scales) might then be most aptly interpreted as changing sensory input to a diffuse, dark, and generally unremarkable stimulus image rather than disconnecting it altogether. Still, visual mental imagery is more likely to be reported when eyes are closed rather than open (Sulfaro et al., 2023). However, while eye-closure tends to aid the recall of fine-grained sensory information, potentially by improving mental imagery, recall is not significantly improved compared to merely looking at blank space (Vredeveldt et al., 2011). Yet, we acknowledge that our conceptualisation of eye-closure may be a simplification given that neural activity patterns between eyes-closed and eyes-open states can differ substantially even when both occur in darkness (Marx et al., 2003, 2004). Even so, an eyes-closed state could still improve mental imagery simply due to the signal-to-noise ratio advantage gained by pitting an imagined stimulus representation against a homogenous closed-eye scene, with the additional benefit of hiding attentional distractors.

### Serotonin and acetylcholine: neurologically-plausible bottom-up and top-down weighting agents

This model implies that neural agents exist which modulate the relative contribution of top-down and bottom-up processes. Given that this model predicts that weighting schemes affect veridical and hallucinatory perception as well as mental imagery, agents which modulate these processes might also be apt candidate weighting agents in our current model.

Serotonin may be one such agent that modulates the contribution of bottom-up information during visual experience. Classic hallucinogens act on the serotonergic system via 5HT2A receptors (Nichols, 2016), and cortical serotonin levels are dependent on plasma levels of tryptophan, serotonin’s blood-brain-barrier-permeable precursor (Fernstrom & Wurtman, 1971). Plasma tryptophan increases over the course of the day, peaking in the late evening (Rao et al., 1994), approximately corresponding to the most common time of hallucination onset for individuals with Charles Bonnet Syndrome, peduncular hallucinosis, and Parkinson’s Disease, including for those without any light perception ability (Manford & Andermann, 1998). Further, 5HT2A receptor expression in the ventral visual pathway of Parkinson’s Disease hallucinators is elevated compared to non-hallucinators (Ballanger et al., 2010; Huot et al., 2010), while inverse agonists for this receptor are used to treat Parkinson’s Disease hallucinations (Yasue et al., 2016). Outside of hallucinatory states, serotonin administered in the primary visual cortex of awake macaques acts as a gain modulator, dampening cell responses to external sensory input without affecting stimulus tuning or selectivity profiles (Seillier et al., 2017). In mice, this dampening in V1 was mediated by a 5HT2A receptor agonist (Michaiel et al., 2019).

Additionally, acetylcholine may modulate the contribution of top-down information during visual experience. The ratio of acetylcholine to serotonin is a factor in hallucination aetiology (Manford & Andermann, 1998), and both serotonin and acetylcholine target layer-five cortical pyramidal neurons, cells which have been proposed as essential for manifesting conscious sensory perception (Aru et al., 2019). High concentrations of acetylcholine can transition the activity of layer-five cortical pyramidal neurons from being driven by feedforward inputs to feedback inputs, which may mediate the hallucinatory experience of dreaming (Aru et al., 2020). The concentration of cortical acetylcholine peaks during rapid-eye-movement sleep, where dreaming is most likely, higher than during wakefulness and higher still than during non-rapid-eye-movement sleep where consciousness is greatly impaired (M. G. Lee et al., 2005). Hence, cortical acetylcholine concentrations may modulate feedback connectivity in sensory cortices.

Given their actions in the visual system, serotonin and acetylcholine levels could be plausible factors accounting for contextual and inter-individual differences in the sensory experiences of mental imagery. Accordingly, they could also be considered as factors which modulate inter-layer weightings within our proposed model. However, ascertaining the role of such neurotransmitters in mental imagery research is challenged by a reliance on human models for self-report, indistinct measurement constructs (e.g. vividness ratings), and the restricted access of psychoactive serotonergic drugs. Ultimately, there are many mechanisms by which our model can facilitate hallucinations and such mechanisms may work in tandem. Note that in all cases, our model does not explain how the initial internally-generated source stimulus might be produced, only how it may be represented, regardless of whether it was generated as part of a voluntary thought, involuntary hallucination, or otherwise.

### Mental imagery as a dynamic process

So far, our predictions have been based on simulations where imagery, once generated, continues to be generated indefinitely and without interruption. Under these conditions, internally-generated content can continue to spread downwards until it reaches an equilibrium point and can spread no further. Yet, if imagery is interrupted or generated intermittently, it may never be generated for a period long enough for it to spread to its maximum possible extent, even if the balance of top-down and bottom-up processes favours a very low equilibrium point (i.e. highly-sensory mental imagery). Analogously, even though the maximum possible speed of a sportscar may be faster than that of a van, the sportscar will be outpaced if the van driver keeps the pedal floored while the sportscar driver only taps the pedal on and off. If mental image content can be generated continuously, or supported by a stream of simulated sensory information at lower levels, then the chances of that content spreading to its maximum possible extent are increased.

### Visual versus auditory mental imagery

Though we have focused on visual mental imagery, mental imagery can occur in other sensory modalities, presumably through similar generative feedback. A complete account of mental imagery should be able to predict and explain modality-specific differences in imagined experience. Auditory hallucinations are roughly twice as common as visual hallucinations in schizophrenia and bipolar disorder (Waters et al., 2014), although how experiences differ between modalities specifically for mental imagery is not well understood. Comparisons generally rely on mental image vividness ratings (e.g. Andrade et al., 2014; Betts, 1910; Gissurarson, 1992; Schifferstein, 2009; Sheehan, 1967; Switras, 1978; Talamini et al., 2022). Yet, as such rating systems are ambiguously defined and difficult to meaningfully interpret (Richardson, 1988), findings have been understandably mixed. Here we make predictions about how the quality of mental imagery may differ between visual and auditory mental imagery, perhaps the two most-studied imagery modalities, based on differences in the brain’s ability to synthesise sensory information in each modality.

Unlike with visual stimuli, our bodies are readily capable of producing their own auditory stimuli, using speech. Self-initiated movements, including the act of speech, produce signals (e.g. efference copies) which carry temporally-precise predictions of the sensory consequences of such movements (Curio et al., 2000). Sensory predictions related to motor movements have been suggested to aid mental imagery when they are produced in the absence of real movements (Gelding et al., 2019; Jack et al., 2019; Pruitt et al., 2019; Scott, 2013; Whitford et al., 2017). This could occur by supplying imagination with the sensory content needed to generate a mental image or by supporting existing mental imagery with a stream of sensory predictions linked to real or simulated movements. As discussed previously, anchoring mental images to continuous sensory information should improve the quality of mental imagery under our model.

While visual mental images could be supported by sensory predictions from real or simulated eye movements, eye movements generally contain information about *where* imagined content is, was or will be (e.g. Fourtassi et al., 2017). However, they are far less informative about *what* is actually being imagined in a given region of the visual field. In comparison, consider inner speech, a type of auditory imagery specifically relating to the imagination of oral sound content like words. If auditory imagery can be expressed as inner speech, such as through converting imagined sounds to onomatopoeia, then real or simulated vocal movements could produce a stream of temporally-precise sensory information not just informing *when* a particular imagined sound should be heard, but also *what* the imagined sound content should be. In essence, using motor systems, the brain could have a greater capacity to simulate sensory features of imagined sounds compared to images, provided the imagined sounds are somewhat possible to replicate vocally. Yet, even when not readily imitable, sounds could still be imagined as some form of onomatopoeia, sacrificing real-world accuracy in order to boost the sensory quality of the imagery experience. Consequently, we expect that auditory mental imagery may often be a more compelling sensory experience than visual mental imagery.

### Limitations and future directions

The primary intention of this article is to highlight that competition from externally-generated sensory input could be a major factor which explains the unique phenomenology of mental imagery. Our modelling is mainly intended merely to provide a simple example of how treating perception as a competitive process can explain aspects of imagery, supporting the larger theoretical argument that mental imagery should always be considered in tandem with cooccurring veridical perceptual processes. Nonetheless, both our argument and our suggested model rely on some assumptions.

Our most crucial restraint is that we require that internally- and externally-generated perceptual processes utilise the same neural substrates such that interference can occur. Our current understanding is that these processes do overlap (Dijkstra et al., 2017, 2020; Ffytche et al., 1998), down to the laminar cortical (Bergmann et al., 2019; Lawrence et al., 2018), if not cellular (Aru et al., 2020), level. However, differences between each process may mean that overlap might vary across the visual hierarchy (e.g. Lawrence et al., 2018), influencing the degree or nature of interference. Yet, even if these processes do overlap, imagined representations may not necessarily propagate smoothly along a serial, retrohierarchical cascade. If imagery utilises direct feedback connections from associative cortices to primary sensory cortices, rather than via a series of intermediate cortical units, interference may have radically different outcomes.

We also rely on assumptions about how neural activity translates into subjective experience. Namely, we assume that lower and higher levels of processing respectively contribute to sensory and abstract features of perceptual experience. Yet, encoding along the visual hierarchy may only correlate with the degree to which semantic content of a perceptual experience is simple or complex, rather than the degree to which such content is experienced in a sensory and modality-specific, rather than non-sensory, way at all. Of course, ascertaining the neural substrates of subjective experience is an immense area of investigation.

Although we make broad predictions about the decaying influence of internally-generated stimuli towards primary sensory regions, the exact balance of this influence, and its dynamics, would likely vary substantially with more neurologically-plausible methods of simulating interference beyond the simple weighted-averaging rule used to approximate interference in our model. Note also that our model does not comment on any of the nonlinear transformations necessary to encode a representation within a layer, but only makes assumptions about how such representations, once formed, may be transmitted and combined. Modelling a more neurologically-plausible system would be a logical next step for investigating mental imagery as a competitive process. Further, it is entirely plausible that other mechanisms aside from competition could account for the segregated distributions of internally- and externally-generated content in the visual system. We merely show that simple models of competition can account for these distributions and many other imagery phenomena.

Our simulations also use retinal inputs (and generally imagined stimulus inputs) which are static and constant, so our simulations cannot account for the effects of neural adaptation on perceptual interference. In reality, retinally-stabilised images tend to induce a reversible blindness, greying-out the visual field. However, such blindness is suspected to be a very early, low-level effect, at or near the retina (Martinez-Conde et al., 2004). Within our model, this state should then be roughly equivalent to feeding in a homogenous field as a real stimulus input, such that the same logic regarding the imagery improvements (or lack thereof) associated with eye-closure and the perception of a dark room would also apply to the perceptual greying-out that occurs with retinally-stabilised images. However, neural adaptation could have many more consequential effects on mental imagery which future studies could explore.

Our model also assumes that feedforward and feedback processes are computationally equivalent. However, Koenig-Robert & Pearson (2021) argue that while feedforward signals initiated from visual input may drive action potentials in the visual cortex, feedback signals terminating in early visual areas tend to be more modulatory, guiding but not overriding activity induced by sensory input. They suggest that this asymmetry could account for the apparent difference in vividness between veridical and mental imagery. However, weak feedback could also be a logical consequence of competition within our model. Our simulations predict that the influence of internally-generated stimulus information should gradually diminish towards lower levels in the presence of bottom-up competition. At some point, this declining influence could fall below a threshold such that it can no longer drive action potentials. Yet, our model still uniquely predicts that feedback should remain influential when external competition is not present. Evidence supports this prediction, as feedback can drive action potentials in the visual cortex of neurotypical individuals during dreaming (Aru et al., 2020), including in neurons which are arguably crucial for manifesting conscious sensory perception (Aru et al., 2019). Hence, it’s plausible that weak feedback follows from perceptual competition. Our model may then supplement the account of Koenig-Robert & Pearson (2021) which could otherwise be interpreted as precluding the possibility of purely top-down hallucinations. Asymmetries in feedforward and feedback processing can still be operationalised in our model by manipulating bottom-up and top-down weightings, but our model distinctly predicts that imagined percepts should be experienced in a less sensory manner than veridical percepts, even if feedforward and feedback processes are computationally equivalent, due to the interference caused by competing sensory input.

Overall, vindicating or falsifying competition as the origin of the quasi-sensory experience of mental imagery will ultimately require a detailed understanding of how neural representations of internally- and externally-generated perceptual content interact. Future studies may seek to explore the predicted impact of hierarchical competition on neural activity using alternative models of feedforward-feedback interference, although our model still capably emulates findings on how imagined and veridical information is distributed in real human brains (e.g. Lee et al., 2012; Spagna et al., 2021; VanRullen & Reddy, 2019).

## Conclusion

Overall, this work provides a formal framework for explaining the unique quasi-sensory experience of mental imagery. We propose that internally-generated thoughts are experienced along an axis from highly-abstract to highly-sensory in nature, aligning with the degree of dimensionality that a given thought is represented in. Under this assumption, we show that competition from bottom-up sensory input may prevent imagined stimuli from being perceived in any substantially sensory way. Accounting for competition in a hierarchical system can therefore provide sufficient conditions for distinguishing imagined, veridical, and hallucinatory perception, as well as variation within these phenomena. As modelling imagery competitively may resolve major lingering questions in imagery research, we recommend future studies investigate the exact manner by which internally- and externally-generated stimuli may be combined in the brain. Ultimately, we conclude that mental imagery is most logically understood as an intrinsically competitive process and should be largely considered as such in ongoing research.

## Acknowledgements

Alexander is supported by an Australian Government Research Training Program Scholarship. Amanda is supported by an Australian Research Council Discovery Early Career Researcher Award (DE200101159). Thomas is supported by an Australian Research Council Discovery Project (DP200101787).

## Data availability

No data was collected as part of this study.

